# Lysineless HiBiT and NanoLuc Tagging Systems as Alternative Tools Monitoring Targeted Protein Degradation

**DOI:** 10.1101/2024.05.14.594249

**Authors:** Hanfeng Lin, Kristin Riching, May Poh Lai, Dong Lu, Ran Cheng, Xiaoli Qi, Jin Wang

**Affiliations:** The Verna and Marrs McLean Department of Biochemistry and Molecular Pharmacology, Baylor College of Medicine, Houston, Texas 77030, United States; Promega Corporation, 2800 Woods Hollow Road, Madison, Wisconsin 53711, United States; Malvern Panalytical Inc., 2400 Computer Drive, Westborough, Massachusetts 01581, United States; Department of Molecular and Cellular Biology, Baylor College of Medicine, Houston, Texas 77030, United States

## Abstract

Target protein degradation (TPD) has emerged as a revolutionary approach in drug discovery, leveraging the cell’s intrinsic machinery to selectively degrade disease-associated proteins. Proteolysis-Targeting Chimeras (PROTACs) exemplify this strategy, exploiting heterobifunctional molecules to induce ubiquitination and subsequent degradation of target proteins. The clinical advancement of PROTACs underscores their potential in therapeutic intervention, with numerous projects progressing through clinical stages. However, monitoring subtle changes in protein abundance induced by TPD molecules demands highly sensitive assays. Nano-luciferase (nLuc) fusion proteins, or the NanoBiT technology derived from it, offer a robust screening platform due to their high sensitivity and stability. Despite these advantages, concerns have arisen regarding potential degradation artifacts introduced by tagging systems due to the presence of lysine residues on them, prompting the development of alternative tools. In this study, we introduce HiBiT-RR and nLuc^K0^, variants devoid of lysine residues, to mitigate such artifacts. Our findings demonstrate that HiBiT-RR maintains similar sensitivity and binding affinity with the original HiBiT. Moreover, the comparison between nLuc^WT^ and nLuc^K0^ constructs reveals variations in degradation patterns induced by certain PROTAC molecules, emphasizing the importance of choosing appropriate tagging systems to ensure the reliability of experimental outcomes in studying protein degradation processes.

## Introduction

Targeted protein degradation (TPD) is a groundbreaking approach in drug discovery that utilizes intrinsic machinery of the cell to selectively degrade disease-associated proteins. Unlike traditional therapeutics that inhibit protein function, TPD molecules (also known as degraders) modulate the protein levels within the cell, providing a unique strategy to tackle diseases. A key player in the TPD field is Proteolysis-Targeting Chimeras (PROTACs). PROTACs are heterobifunctional molecules that can provide body-wide phenocopy knock-down without modification on the genome. The warhead part can bind to the target protein and the E3 ligand part can bind to E3 ubiquitin ligase to facilitate the formation of a ternary complex, leading to ubiquitination on surface-exposed lysine residues of the target protein, ultimately leading to targeted protein degradation^1,2^. The ability of PROTACs to harness the natural protein degradation pathway has broadened the scope of “druggable” targets, offering new possibilities for therapeutic intervention. As of 2023, a total of 26 PROTAC projects worldwide have entered the clinical stage. The most rapid progression was observed in the ARV-471 project, a collaborative effort between Arvinas and Pfizer, which commenced phase III clinical trials in 2022^3–6^. Simultaneously, several other degraders are either already in the initial stages of clinical trials or are poised to enter them. The PROTAC field is flourishing, with a multitude of degraders competing for a place in the clinical market.

To monitor the subtle target protein abundance change induced by the TPD molecules in the complex cellular environment, a highly sensitive and high-throughput compatible protein abundance assay with broad dynamic range is ideal for the screening purpose. Nano-luciferase (nLuc), a small 19-kDa, highly stable, ATP-independent, bioluminescent protein engineered from the original deep-sea shrimp luciferase (Oplophorus-luciferin 2-monooxygenase), has been utilized to develop robust and ultra-high sensitivity screening systems^7^. As a commonly used chemical biology tool in the high throughput evaluation of degrader efficacy, strong bioluminescence signal generated from the nLuc fusion protein could sensitively reflect real-time fusion protein level in live cells or crude cell lysates, allowing for the kinetic monitoring of protein degradation event upon degrader treatment^8,9^. To further minimize the tagging size, NanoBiT technology was utilized^10^. This approach involves a complementing system where an 11-amino acid peptide (HiBiT) is tagged onto the protein of interest, enabling interaction with an 18 kDa polypeptide (LgBiT). This interaction results in the formation of an active luciferase capable of producing light upon reaction with its luminogenic substrate, furimazine or its stabilized version endurazine^8,10,11^.

The efficient ubiquitination of a substrate lysine requires the appropriate spatial proximity between the substrate lysine and the narrow catalytic site on the E2 enzyme^12–14^. This spatial arrangement is crucial for facilitating the ubiquitination reaction. Therefore, the accessibility and positioning of lysine residues play a critical role in determining their susceptibility to ubiquitination^15,16^. It has been reported that lysine residues of the target protein located on the ubiquitin accessible band of the E3 ligase machinery will contribute more to the degradation^16–18^. Therefore, recently there has been increasing concerns about the introduced lysine residues on the tagging protein or peptide may contribute to degradation artifacts. A typical example was the successful degradation of GFP-KRAS^G12C^ but not the endogenous KRAS^G12C^ by XY-4-88^19^. For those target proteins with only a few lysine residues on the surface, the artificially introduced lysine on the tag may even mislead the researchers for pursuing a non-degradable PROTAC molecule.

In this study, we present two alternative tools, HiBiT-RR and nLuc^K0^, in which all the lysine residues have been replaced by arginine residues from their original sequences, to avoid potential risk from degradation artifacts. We show here that HiBiT-RR shows a comparable binding affinity towards LgBiT to the original HiBiT peptide, with no sacrifice on the luminescence intensity. HiBiT-RR tag is as sensitive as HiBiT in detecting and quantifying drug-induced protein degradation. On the other hand, by comparing nLuc^WT^ and nLuc^K0^ construct, we report that some PROTAC molecules tested in this paper can trigger stronger degradation in the nLuc^WT^ fusion protein than the nLuc^K0^ counterpart. This discovery suggests that degradation artifacts stemming from the tagging system might be more widespread than currently understood. It also highlights the importance of choosing the right tagging system to help minimize potential interference or artifacts in studying protein degradation processes, thereby enhancing the reliability and validity of experimental results.

## Results

### HiBiT-RR showed comparable LgBiT protein binding affinity with HiBiT-KK

Lysine is the most common residue to accept ubiquitin transfer from E2 enzymes due to their reactivity. In order to deal with the potential artificial degradation of the target protein originated from the two lysine residues in the original version of HiBiT tag (VSGWRLF<u>KK</u>IS), referred as HiBiT-KK thereafter ^10,11^, we started to characterize the biophysical and enzymatic properties of the lysine-less version of the HiBiT (VSGWRLF<u>RR</u>IS), hereinafter referred as HiBiT-RR, by substituting the two lysine with arginine residues (**Figure 1A**). We first characterized the binding between LgBiT protein and HiBiT-KK or HiBiT-RR. Fixing the amount of LgBiT, we titrated a series of HiBiT concentrations for both KK and RR variants. Both HiBiT-KK and HiBiT-RR showed low nM EC_50_ values and comparable luminescence signal intensities no matter whether our homemade His-LgBiT protein or the LgBiT protein from Promega Corp. was used (**Figure 1B,C**). A comparable level of luminescence signal is as expected since the KK to RR mutation site is distant from the luciferase catalytic pocket according to a recently deposited nLuc structure (PDB: 7SNT). However, the EC_50_ value we got from LgBiT-HiBiT interaction was ∼ 10 times weaker than previously reported 700 pM^11^. Therefore, we performed a reverse titration of various concentrations of LgBiT protein into a fixed amount of HiBiT variants. After subtracting the signal from LgBiT itself in the absence of any HiBiT peptide, both HiBiT-KK or RR still showed similar EC_50_ levels as are in the forward titration (**Figure 1D,E**). The EC_50_ levels remain unchanged for up to 6 hours incubation, indicating the establishment of equilibrium (**Figure S1**).

**Figure 1.**
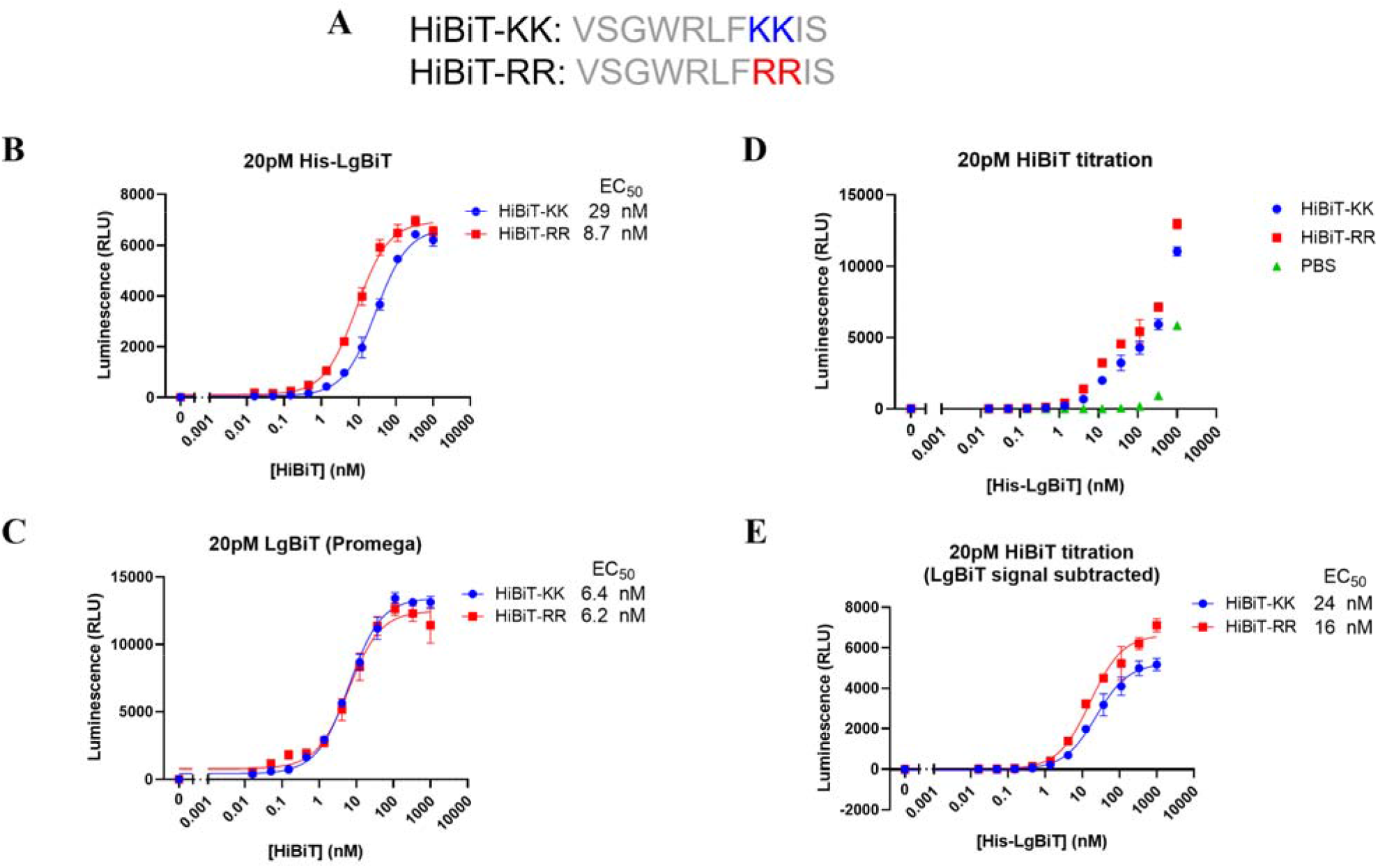
HiBiT-RR showed comparable LgBiT protein binding affinity and similar luminescence output with the original HiBiT-KK. **A**. Peptide sequences of the original HiBiT peptide (HiBiT-KK in this paper) from Promega and the lysine-less version (HiBiT-RR) **B, C**. The luminescence signal from 20 pM LgBiT protein (homemade His-LgBiT for **B** and Promega commercially available LgBiT protein for **C**) when titrating with various concentrations of HiBiT-KK or HiBiT-RR for 30 min. **D**. The luminescence signal from 20 pM HiBiT variants or PBS when titrating with various concentrations of His-LgBiT recombinant protein for 30 min. **E**. The background LgBiT luminescence from PBS group were subtracted from panel D.

We also biophysically determined their affinity and binding kinetics in the absence of enzyme substrate using Grating-Coupled Interferometry (GCI). In parallel with enzymatic measurements, both HiBiT-KK and RR demonstrated an affinity at the single-digit nanomolar level towards His-LgBiT. Notably, HiBiT-RR exhibited a marginally superior performance compared to HiBiT-KK, displaying a slower dissociation rate (k_off_) under conditions of both high and low density of LgBiT (**Figure 2**).

**Figure 2.**
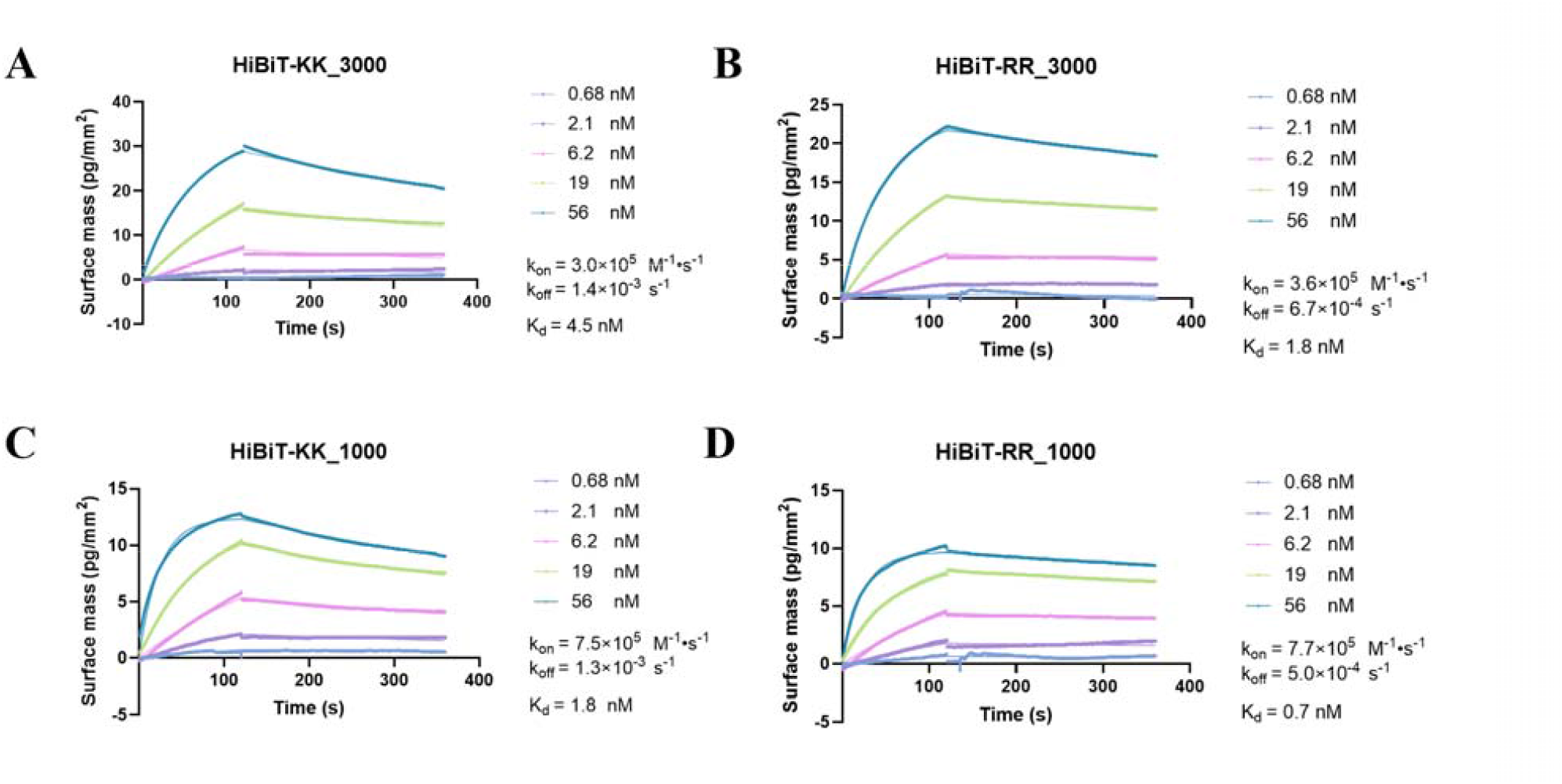
Grating Coupled Interferometry measurement of HiBiT-KK and RR showed comparable LgBiT protein binding kinetics. Various concentrations of HiBiT-KK (**A, C**) or HiBiT-RR (**B, D**) were flow against immobilized His-LgBiT at high (**A, B**) or low (**C, D**) surface density. The association and dissociation events were monitored real-time using Grating-Coupled Interferometry.

All these evidence indicates that under such experimental condition, the binding affinity between HiBiT and LgBiT in the presence of enzyme substrate may not be as tight as determined before in sub-nM range, yet still tight enough for applications such as sensitively detecting HiBiT in cell lysates. Also, introducing the RR mutation into the established HiBiT sequence does not compromise the readout intensity, from which a robust luminescence assay with good sensitivity can be developed.

### HiBiT-RR tagged protein degradation induced by degraders

Protein degrader molecules like PROTACs or molecular glue are gathering increasing attention as both therapeutic modalities and chemical biology tools. The warhead component of these molecules binds to the target protein, while the E3 ligand portion interacts with the E3 ubiquitin ligase, facilitating the formation of a ternary complex. This complex, in turn, triggers ubiquitination, leading to the eventual degradation of the target protein. A pivotal step in the protein degradation pathway is the transfer of ubiquitin onto lysine residues exposed on the protein surface, navigating the protein to the proteasome for degradation. There have been concerns since the release of HiBiT-tagging system about the existing lysine in the HiBiT sequence, with apprehensions about whether these artificially introduced lysine residues could potentially contribute to the degradation of the target protein. This concern becomes particularly relevant in the case of protein degraders since the protein of interest has already been recruited and loaded onto the E3 ligase complex.

Here we characterized the degradation efficiency of four BTK PROTACs (RC-1^20^, NX-2127^21,22^, NRX-0492^23^ and DD-03-171^24^) covering different target engagement mechanisms (RC-1 for reversible covalent PROTAC, others for reversible PROTAC) using BTK kinase domain (residue 382-659, BTK_KD_) tagged with either HiBiT-KK or RR. (**Figure 3**). Results show that the HiBiT-RR construct (**Figure 3B, D**) was able to reproduce those characteristic parameters of PROTACs observed in the traditional HiBiT-KK construct (**Figure 3A, C**) such as DC_50_, D_max_, as well as the strong hook effect observed in RC-1 at high doses. In our previous research, RC-1 did not show a very strong hook effect, we believe it was due to the difference of the cooperativity between BTK_KD_ and full length BTK. Indeed, in Ramos BTK-HiBiT-KK knock-in cell line, we observed a weaker hook effect and a deeper D_max_ (**Figure S2**).

**Figure 3.**
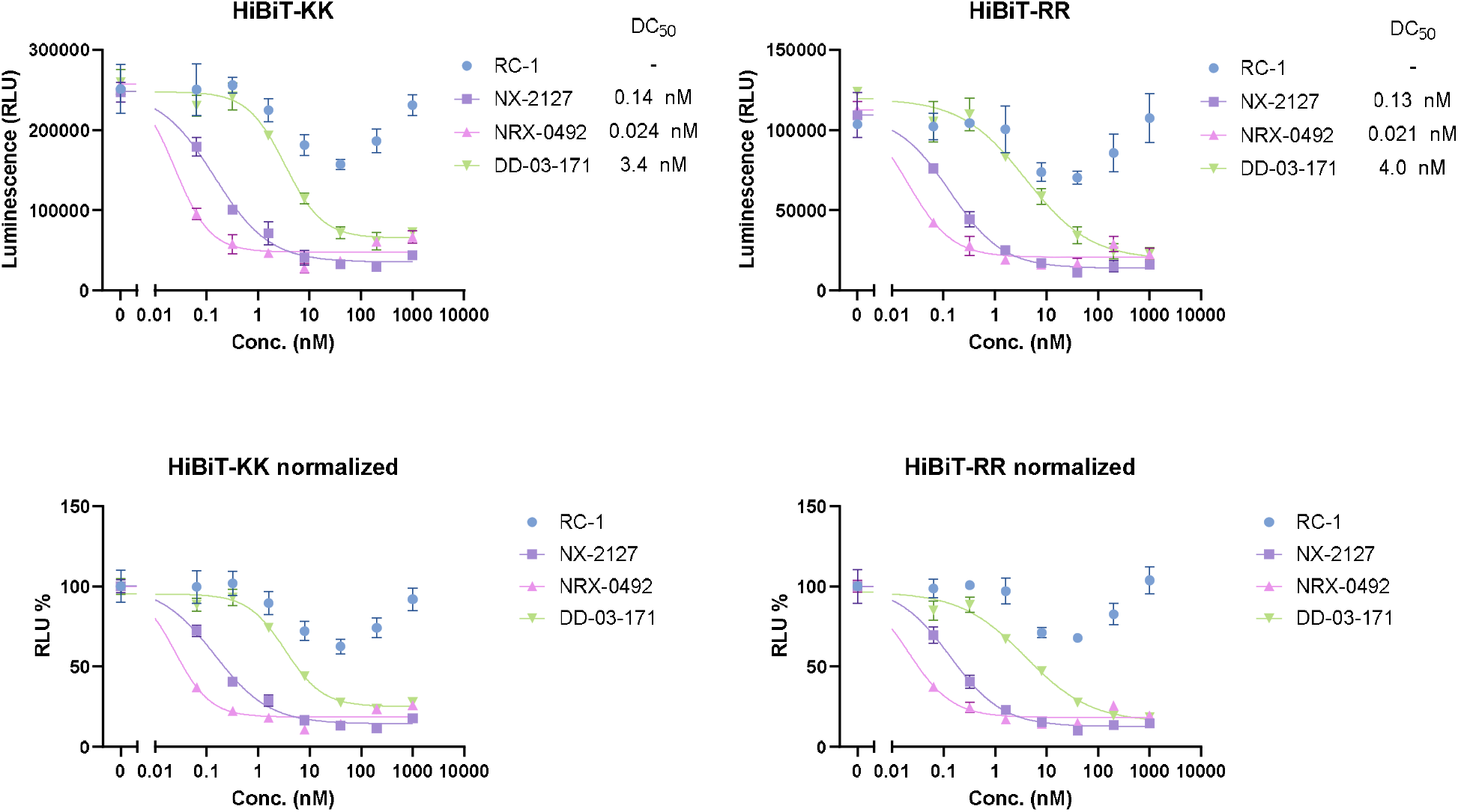
Comparison of PROTAC degradation potency of HiBiT-KK or HiBiT-RR tagged BTK kinase domain in HEK293T cell line. HEK293T cells transiently transfected with BTK_KD_-HiBiT-KK **(A)** or BTK_KD_-HiBiT-RR **(B)** were transferred into 96-well plate for 20,000 cells per well. Cells were incubated with DMSO or indicated compounds at 0.064, 0.32, 1.6, 8, 40, 200, 1,000 nM in 1% DMSO for 24 hours. Bioluminescence signal was generated by adding furimazine substrate and LgBiT purified protein in the cell lysis buffer. **C,D**. Bioluminescence signal from A,B normalized to their corresponding DMSO groups.

With the robustness of transiently expressed HiBiT-RR construct confirmed, we tested both systems in the context of HiBiT *in situ* knock-in cell lines. By CRISPR knocking in either HiBiT-KK or -RR at the N-terminus of BRD4 in HEK293 cell line, MZ1 and dBET6, two classical BRD4 PROTACs ^25,26^, were further used to characterize the HiBiT-KK or -RR knock-in cell lines. Similarly, no significant DC_50_ or D_max_ difference was observed for MZ1 degrading HiBiT-BRD4 (**Figure 4A-D**). For dBET6, we observe marginal difference that HiBiT-RR BRD4 undergoes a slower degradation kinetics than HiBiT-KK in high concentration dBET6 treatment, although a similar D_max_ was observed, and the degradation potency when plateau was achieved shows less than 2-fold difference (**Figure 4E-H**). Altogether, our data indicates that HiBiT-RR also works as well as the original HiBiT-KK in the *in situ* genome knock-in systems.

**Figure 4.**
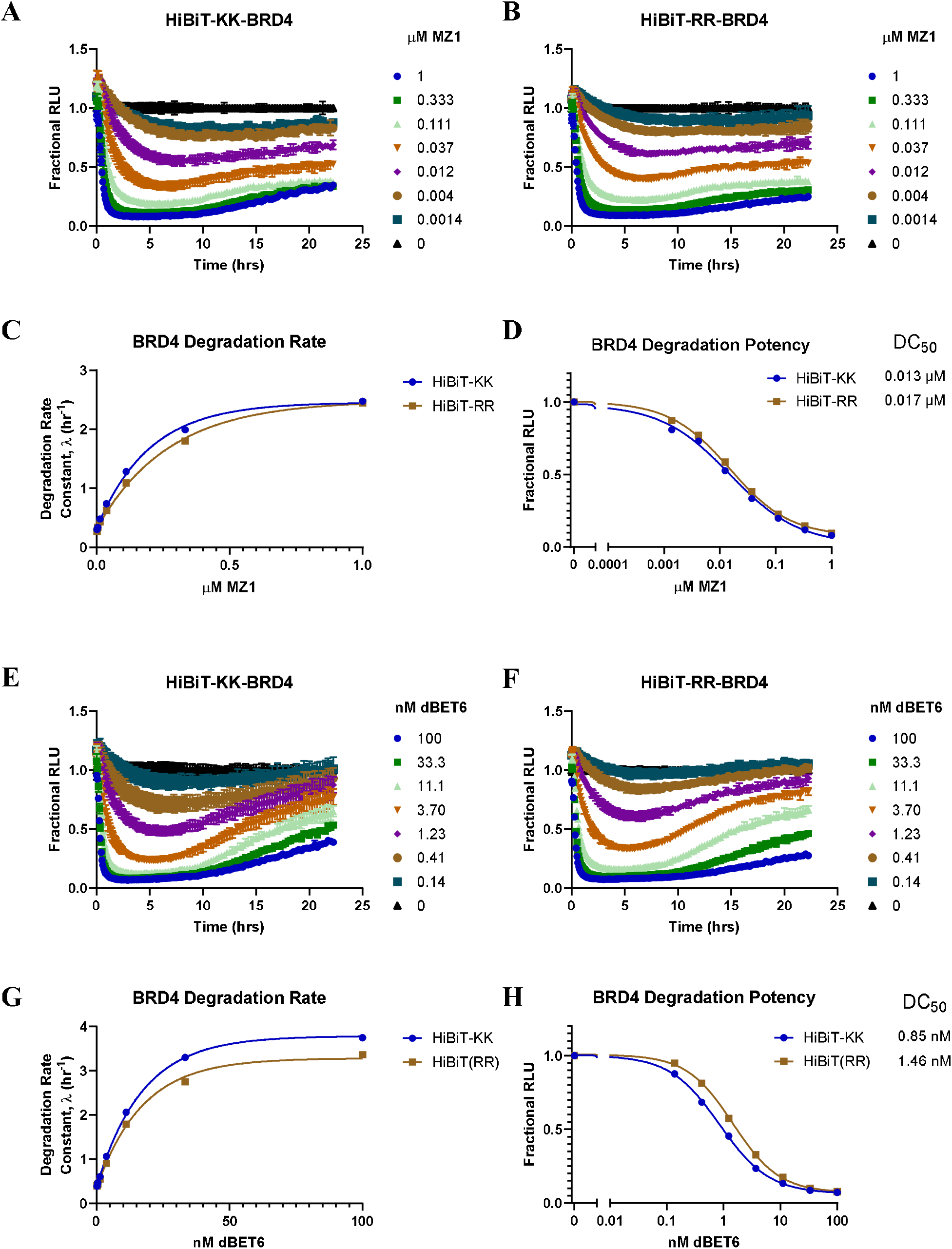
A HiBiT-KK CRISPR knock-in BRD4 in HEK293 cell line was compared to HiBiT-RR knock-in using dBET6 (A-D) and MZ1 (E-H). **A-B, E-F**. Degradation kinetics of HiBiT-KK or -RR BRD4 normalized to DMSO group. **C, G**. HiBiT-KK or RR BRD4 degradation rate of dBET6 or MZ1 fitting from the initial phase (before 7 hours) **D, H**. HiBiT-KK or RR BRD4 degradation potency comparison under dBET6 or MZ1 treatment when plateau was achieved.

### Exploring the potential of nLuc^K0^ tagged protein in targeted protein degradation

So far, we have validated HiBiT-RR as a feasible tool for protein abundance monitoring, which shows comparable sensitivity with HiBiT-KK but ablated the potential pitfall of degradation artifacts. However, using HiBiT-tag protein alone does not allow continuously monitoring the protein level changes in the time course study, unless an additional copy of LgBiT was also knocked into the genome^8^. However, potential ubiquitination could still happen on the lysine residue of the wildtype LgBiT. Ectopically expressing a nLuc fusion protein is a common way to evaluate degradation kinetics. Here we would also like to report our attempt of using a no-lysine version of nLuc (nLuc^K0^), where all the lysine residues in the original nLuc sequence were mutated into arginine (sequence as shown in **Figure 5A**). By fusing nLuc WT or K0 to the N-terminus of RIPK1 kinase domain (RIPK1_KD_) and expressing it in HEK293T cells, we head-to-head compared the degradation capacity of a series of RIPK1 degraders (**Table S1**) against these constructs (**Figure 5B, C**). For majority compounds, both nLuc WT and K0 construct show similar maximal degradation (D_max_) and DC_50_ values, while the negative control compound 4172NC shows no degradation in both constructs. But for compound 5037, we observed a 3-fold worsening in DC_50_ when nLuc^K0^ construct was used (nLuc^WT^ 47 nM versus nLuc^K0^ 147 nM). A similar potency reduction was also observed in compound 5077, a CRBN-based RIPK1 PROTAC (nLuc^WT^ 10 nM versus nLuc^K0^ 39 nM). This observed potency reduction could be due to the removal of the ubiquitin-able lysine residues on the nLuc surface, highlighting the potential degradation artifacts when ubiquitination happens on the nLuc protein. Transition to the K0 construct could avoid such degradation artifacts from the nLuc part. Immunoblot confirmed the degradation of both nLuc-RIPK1_KD_ fusion protein (**Figure 5D**).

**Figure 5.**
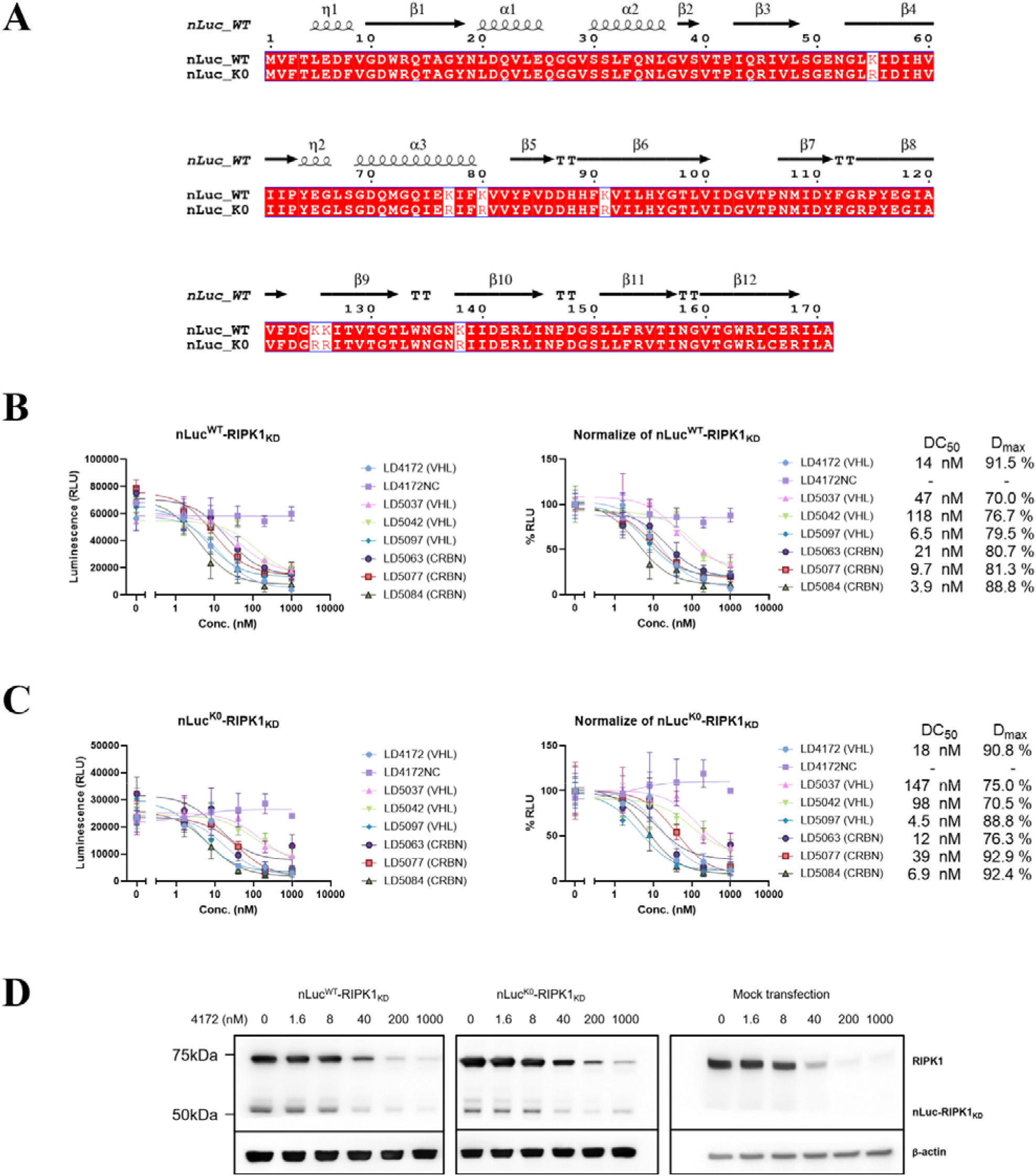
Comparison of nLuc^WT^ and nLuc^K0^ tagged RIPK1 kinase domain degradation potency in transfected HEK293T cell line. **A**. Sequence comparison between nLuc^WT^ and nLuc^K0^. Secondary structure information extracted from PDB 5IBO. **B, C**. HEK293T cells transiently transfected with nLuc^WT^-RIPK1_KD_ (B) or nLuc^K0^-RIPK1_KD_ (C) were transferred into 96-well plate and incubated with DMSO or indicated compounds at 1.6, 8, 40, 200, 1,000 nM in 1% DMSO for 24 hours. Bioluminescence signals were recorded in live cells using furimazine-containing OptiMEM. Unit for Bottom and Top: %, IC_50_: nM. **D**. Immunoblot of RIPK1 in HEK293T cells transfected with nLuc^WT^/^K0^-RIPK1_KD_ plasmid or empty vector after 24h treatment of 4172.

Since we observe a lower luminescence intensity when expressing the nLuc^K0^ construct in the mammalian cell, we purified both His_6_-tagged nLuc WT and K0 protein from *E. coli* BL21(DE3) (**Figure S3A, B**). Surprisingly, we found that nLuc^K0^ protein has significantly decreased enzyme activity (up to 400-fold) compared to nLuc^WT^ under the same *in vitro* assay condition (**Figure S3C**). This result could explain the decrease luminescence intensity from nLuc^K0^ when overexpressed in mammalian cells. nLuc^K0^ protein activity may be heavily rely on the cellular environment to maintain the protein stability.

## Discussion

Target Protein Degradation (TPD) represents a revolutionary approach in drug discovery, harnessing the intrinsic machinery of cell protein degradation pathway to selectively degrade disease-associated proteins. Unlike conventional therapeutics that inhibit protein function, TPD molecules, also known as degraders, modulate protein levels within cells, offering a unique strategy to combat diseases. However, amidst this progress, there exist challenges in accurately monitoring subtle changes in target protein abundance induced by TPD molecules within complex cellular environments. Immunoblot has long been used as the golden standard to detect and quantify target protein degradation, but the long processing steps, the limited numbers of samples in each gel make the throughput very low, sometimes even lead to artifacts. Capillary electrophoresis immunoassay typically requires less sample consumption, involves simpler procedures, and shortens analysis time^27^. It has been applied to monitor BTK level changes^28^, and BRD4 bromodomain ubiquitination level^29^ after PROTAC treatment. Time-Resolved Fluorescence Resonance Energy Transfer (TR-FRET) has recently been developed to quantify the protein level in the crude lysate in a high-throughput manner^30^. However, all these methods depend on highly specific antibody-antigen interaction. In addition, the TR-FRET based assay also requires development of tracer compounds. To address these problems, highly sensitive and high-throughput compatible protein abundance assays with broad dynamic ranges are indispensable for screening purposes.

Reporter assays represent a rapid and sensitive method to measure protein degradation based on fluorescence or bioluminescence. GFP has been used to quantitatively monitor the target protein degradation event for KRAS^G12C 19^ and BRD4 bromodomains^26^. The development of a complementary luciferase based HiBiT tagging system offered another reporter assay to monitor protein degradation. The high affinity between HiBiT and LgBiT complex allows immediate quantification of HiBiT tagged protein using bioluminescence readout. This system has been successfully applied in kinetically monitoring the potency of BRD2/3/4 PROTAC MZ1^8^ and BRD7/9 PROTAC VZ185^31^, IMiD molecular glues^32^, kinases^32,33^ and fusion-tag PROTACs^34^. However, there has been reported cases that fusion tags on the target protein may alter the degradability of the target protein by introducing artificial ubiquitination sites^19^.

Since there is currently no way to guarantee whether a fusion tag may contribute to the protein degradability by introducing novel ubiquitination sites, as an expansion of current chemical biology toolbox, this study introduced two alternative tagging systems, HiBiT-RR and nLuc^K0^, aimed at minimizing potential degradation artifacts during TPD molecule discovery processes. HiBiT-RR, a lysine-less version of the HiBiT tag, demonstrated comparable binding affinity to LgBiT protein with the original HiBiT-KK, ensuring reliable protein abundance monitoring without sacrificing assay sensitivity. Moreover, HiBiT-RR maintained similar degradation efficiency as HiBiT-KK when tested with various BTK and BRD4 PROTACs, indicating its suitability for evaluating degrader potency. Interestingly, we observed in the HiBiT-RR BRD4 knock-in cells, only dBET6 showed a slightly slower degradation kinetics compared to HiBiT-KK, but not MZ1. This suggests that the fusion tag may or may not contribute to the degradation event depending on the compound used. It is plausible that the tag on the target protein adopts a conformation that is more favorable for accepting ubiquitin in the presence of certain compounds but not in others. This nuanced understanding underscores the complex interplay among the tagging system, compound-induced ternary complex conformation, and protein degradation mechanism.

Furthermore, the study explored the potential of nLuc^K0^, a no-lysine version of nLuc, in targeted protein degradation studies. While both nLuc^WT^ and nLuc^K0^ constructs exhibited similar degradation potency for most compounds tested, a notable reduction in degradation potency was observed for specific compounds with nLuc^K0^, suggesting potential degradation artifacts stemming from ubiquitination on the lysine residues of nLuc^WT^. Transitioning to the nLuc^K0^ construct could mitigate such artifacts, enhancing the reliability of degradation studies. However, it is worth noting that the relative lower luminescence intensity observed with the nLuc^K0^ construct when expressed in mammalian cells could be attributed to decreased enzyme activity compared to nLuc^WT^. The enzyme activity difference was even more obvious using purified nLuc^WT^ and nLuc^K0^ protein, which indicates the instability nature of nLuc^K0^ in non-cellular environment. But out of all the PROTACs we tested, we did not find any compounds that show more than 2-fold stronger degradation potency in the nLuc^K0^ than nLuc^WT^ construct. This observation helps alleviate concerns regarding potential artificial degradation due to instability of the nLuc^K0^ protein. Although the luminescence signal decrease in cellular assay is trivial and can be easily compensated by increasing instrument photomultiplier tube (PMT) gain setting, this finding underscores the importance of considering the influence on cellular environment when overexpressing proteins with less stability, especially in the context of assay development and interpretation.

Overall, the study underscores the significance of choosing appropriate tagging systems, such as HiBiT-RR and nLuc^K0^, to minimize potential interference and artifacts in studying protein degradation processes. By enhancing the reliability and validity of experimental results, these tools contribute to advancing drug discovery efforts utilizing TPD molecules, ultimately leading to more effective therapeutic interventions.

## Methods

### Reagents & Cell culture

Unless otherwise stated, all chemicals were purchased from Sigma Aldrich. HiBiT-KK and RR peptides were synthesized by GenScript, USA. pcDNA3.1 vector backbone was from GenScript, USA. DH5_α_ and BL21(DE3) *E. coli* strains were from Thermo Scientific (Cat. No. EC0111, EC0114).

HEK293T/17 cell line (hereinafter referred as HEK293T) was purchased from American Type Culture Collection (Cat. No. CRL-11268). HEK293T cells were maintained in DMEM (Corning, Cat. No. 10-013-CV) supplemented with 10% fetal bovine serum (FBS) (Corning, Cat. No. 35-011-CV) at 37°C with 5% CO_2_. Transfections were carried out using the homemade calcium phosphate method. Generally, for a 10 cm dish, 2 million HEK293T cells in 10ml medium were plated the day before transfection. The desired amount of plasmid was mixed with 75 µL 2 M CaCl_2_ and 550 µL sterile H_2_O, then 625 µL 2× HBSS buffer (50 mM HEPES, 280 mM NaCl, 1.42 mM Na_2_HPO_4_, pH 7.05) was added and settled for 15min to allow precipitation formation. The mixture was dropwise added into the culture dish and incubated for 15-18 h. The cells were further incubated in fresh medium for 24-36 h to allow gene expression and then ready for assay purpose.

### Plasmid cloning & Protein purification

nLuc^WT^/^K0^-RIPK1_KD_ (kinase domain 1-324) fusion gene with a 24-residue flexible linker in between (SGGRSSGSGSTSGSGTSLYRRVGT) was synthesized and codon optimized by GenScript and cloned into pcDNA3.1 backbone. For BTK_KD_-HiBiT^KK^/^RR^ plasmids, the BTK-nLuc template was purchased from Promega (Cat. No. N2441). The gene coding for BTK kinase domain (residue 382-659) was subcloned into pcDNA3.1 vector, followed by the site-directed mutagenesis (ClonExpress II One Step Cloning Kit C112, Vazyme) to fuse the HiBiT-KK or HiBiT-RR gene to the C-terminus of BTK kinase domain with a two-residue linker (VS).

Codon-optimized LgBiT insert was cloned into pET-28a(+) vector using NdeI/BamHI restriction sites to obtain pET28a-His_6_-LgBiT plasmid. The plasmid was transformed into BL21(DE3) *E. coli* strain under kanamycin selection pressure. The transformed strain was inoculated into 1 L LB medium and induced by 0.5 mM IPTG when the OD600 value reached 0.8. After shaking the culture overnight at 18 °C, the cells were pelleted and resuspended in the lysis buffer (50 mM Tris-HCl, 150 mM NaCl, 20 mM imidazole, pH 7.5) plus protease inhibitor cocktail pills (Roche). The cells were ruptured by sonication on ice for 30min (4s pulse + 8s rest cycle) with 30% amplitude, followed by 18,000×g 45 min centrifugation at 4°C. The supernatant was mixed and gently rotated for 1h with 5ml Ni-NTA beads (Qiagen) pre-equilibrated with lysis buffer. The beads were further washed using 100 ml lysis buffer and bound protein was eluted by elution buffer containing 50 mM Tris-HCl, 150 mM NaCl, 300 mM imidazole, pH 7.5. Protein purity was checked by SDS-PAGE and Coomassie staining. The elution fraction was buffer exchanged into 50 mM Tris-HCl, 150 mM NaCl, pH 7.5, concentrated to 402 uM, aliquoted and snap frozen for long-term storage at -80 °C. His-nLuc^WT^ and His-nLuc^K0^ was purified using the same protocol.

### Luminescence-based binding affinity titration

For HiBiT titration against LgBiT, the HiBiT-KK or RR peptide was dissolved in ddH_2_O as 1 mM stock solution. LgBiT protein either purified in-house (His-LgBiT) or purchased from Promega (N401A, 200 µM stock) was diluted to 20 pM in PBS pH 7.4 solution and dispensed into white 96 well non-binding surface plate (Corning #3600) with 90 uL/well. HiBiT-KK or RR peptides were 3-fold serial diluted as 10× stock solution and 10 µL was added into each well, giving a HiBiT working concentration ranging from 0.016 nM to 1,000 nM. After equilibrating at room temperature for 30 min, 5 µL 1:25 diluted Nano-Glo luciferase assay substrate (Promega, N113A) in PBS was dispensed into each well and quickly mixed. The luminescence signal was collected using SYNERGY H1 microplate reader (BioTek) with the PMT gain set as 120. EC50 value was calculated by fitting into agonist concentration - response model (Prism 10, GraphPad software, Inc.).

For LgBiT titration against HiBiT-KK or RR, a similar protocol was used except for that 90 µL HiBiT peptides (20 pM) were pre-dispensed into each well, followed by adding 10 µL 3-fold serial diluted LgBiT protein in PBS with a working concentration from 0.016 nM to 1,000 nM. Since high concentration LgBiT protein showed intrinsic catalytic effect even in the absence of HiBiT peptide, we subtracted the baseline luminescence signal of LgBiT-only samples from that of the HiBiT experimental groups and then performed the agonist concentration - response fitting.

### Grating-coupled interferometry (GCI) characterization of peptide-protein binding kinetics

Grating-coupled interferometry experiments were conducted on a Creoptix WAVEdelta system. A PCP-NTA (Polycarboxylate Planar functionalized with Nitriloacetic acid) chip was used to immobilize His_6x_-LgBiT. Specifically, the PCP-NTA was conditioned for 3 mins at 10 ul/min with 0.1 M borate/1M NaCl at pH 9.0 (Xantec Elution buffer). The surface was injected for 2 mins at 10 uL/min with 0.5 mM NiCl_2_. The ligand His-LgBiT was diluted to 50 ug/mL concentration with PBS, then injected directly to the surface at 10 ul/min using target level function. Final surface densities of 3000 pg/mm^2^ (high surface density) and 1000 pg/mm^2^ (low surface density) His-LgBiT were reached on two flow channels, respectively. Traditional Kinetic experiments were performed using running buffer composed of 0.01 M HEPES, 0.15 M NaCl and 0.05% v/v Surfactant P20, pH 7.4. For Traditional Kinetic experiment, various concentrations of peptides HiBiT-KK and HiBiT-RR were injected over the chip surface at flow rate of 30 uL/min with an association time of 120 s followed by 240 s of dissociation. For data analysis, both reference subtraction and blank subtraction were performed to generate the final sensorgrams. The global fitting for kon and koff was achieved through “association then dissociation” non-linear regression model using Prism GraphPad v10.2.

### In vitro protein degradation assay

Two million HEK293T cells were transfected with BTK_KD_-HiBiT^KK^/^RR^ or nLuc^WT^/^K0^-RIPK1_KD_ plasmid in a 10cm dish according to the general transfection protocol as mentioned above. The absolute amount of RIPK1 plasmids used has been optimized through immunoblotting to ensure the expression level does not exceed the endogenous RIPK1 level. The absolute amount of BTK plasmids did not undergo optimization since HEK293T cells do not express BTK at all, but the transfected plasmid amount for BTK_KD_-HiBiT^KK^ and BTK_KD_-HiBiT^RR^ was consistent.

After 24-36 hours expression, cells were trypsinized and resuspended in Opti-MEM I phenol red-free medium (Life Technologies, Cat. No. 11058021) supplied with 4% FBS. 20,000 cells in 99 µL medium were plated into each well of the 96 well plates (Corning, #3917). After settling down for 2-4 hours, cells were incubated with serially diluted BTK PROTAC compounds (from 0.064 to 1000 nM, 5-fold dilution, 1% DMSO) or RIPK1 PROTAC compounds (from 1.6 to 1,000 nM, 5-fold dilution, 1% DMSO) for 24 h at 37 °_C_ with 5% CO_2_ in an incubator.

For nLuc-based assay, we chose to use live cell-based readout to maximize the luminescence signals. The medium was completely aspirated from each well, followed by adding into each well 25 µL Nano-Glo luciferase assay substrate 1:100 freshly diluted in PBS. Luminescence signals were collected using SYNERGY H1 microplate reader (BioTek) after brief shaking.

For HiBiT-based assay, lysis protocol was used to make HiBiT-tagged protein accessible to LgBiT protein. Briefly, Nano-Glo luciferase assay substrate was 1:100 diluted in nLuc lytic buffer ^7^ (100 mM MES pH 6.0, 1 mM CDTA, 0.5% (v/v) Tergitol, 0.05% (v/v) Antiform 204, 150 mM KCl, 1 mM DTT, and 35 mM thiourea) containing 200 nM LgBiT protein to make 2× nLuc substrate lytic buffer. 100 µL 2× buffer was subsequently added into each 96-well. After brief shaking and waiting 5 min for complete cell lysis and equilibration, the luminescence signals were collected.

### Immunoblotting

Transfected HEK293T cells were plated in 12-well plates. After 24h treatment, cells were harvested in 1_x_ RIPA buffer (Thermo Scientific J62524.AE) with 1x Halt™ Protease and Phosphatase Inhibitor Cocktail (Thermo Fisher Scientific, Cat. No. PI78440) and Benzonase Nuclease (Sigma E1014) to reduce sample viscosity. Lysates were centrifuged at 15,000 × g for 10 min at 4 °C and the supernatant was quantified for total protein concentration using the Pierce BCA Protein Assay (Thermo Fisher Scientific, Cat. No. 23225). Total protein 20 ug were separated by SDS-PAGE and further transferred onto 0.2 um PVDF membrane (Millipore, Cat. No. IPVH00010) pre-soaked with ethanol. The membranes were probed overnight at 4 °C with the specified primary antibodies at the dilution of 1:1000 (Cell Signaling Technology: RIPK1 Rabbit mAb#3493)) or 1:4000 (β-actin Rabbit mAb#4970)), followed by the HRP-conjugated secondary antibody (Kindle Biosciences, #R1006, 1:1000) for 1 h at room temperature. Imaging was performed using ultra digital-ECL substrate solution (Kindle Biosciences, #R1002) in the digital imaging system.

### In vitro nLuc enzyme activity assay

His-nLuc^WT^ and His-nLuc^K0^ purified protein were 2-fold serial diluted in 50 mM Tris-HCl, 150 mM NaCl, pH 7.5. A total of 99 µL was dispensed in each well of a 96-well plate. 1µL undiluted Nano-Glo luciferase assay substrate (Promega, N113A) was dispensed into each well and quickly mixed. The luminescence signal was collected using SYNERGY H1 microplate reader (BioTek) with the PMT gain set as 120.

### Data analysis

All the experiment data was from at least n=3 technical replicates and cell-based experiment contains at least two batches of biological replicates. Degrader dose-response data was fitted using inhibitor vs response three parameters model, luminescence-based binding affinity titration data was fitted using agonist vs response three parameters model in GraphPad Prism 10.

## Acknowledgement

This research was supported in part by the National Institute of Health (R01-CA250503 and R01-CA268518 to J.W.), the Cancer Prevention & Research Institute of Texas (CPRIT, RP220480 to J.W.), and Michael E. DeBakey, M.D., Professor in Pharmacology (to J.W.). We also appreciate all suggestions and advice from Dr. Chris Eggers at Promega.

