## Supplemental Info for "Lysineless HiBiT and NanoLuc Tagging Systems as Alternative Tools Monitoring Targeted Protein Degradation"

### Supplementary Material

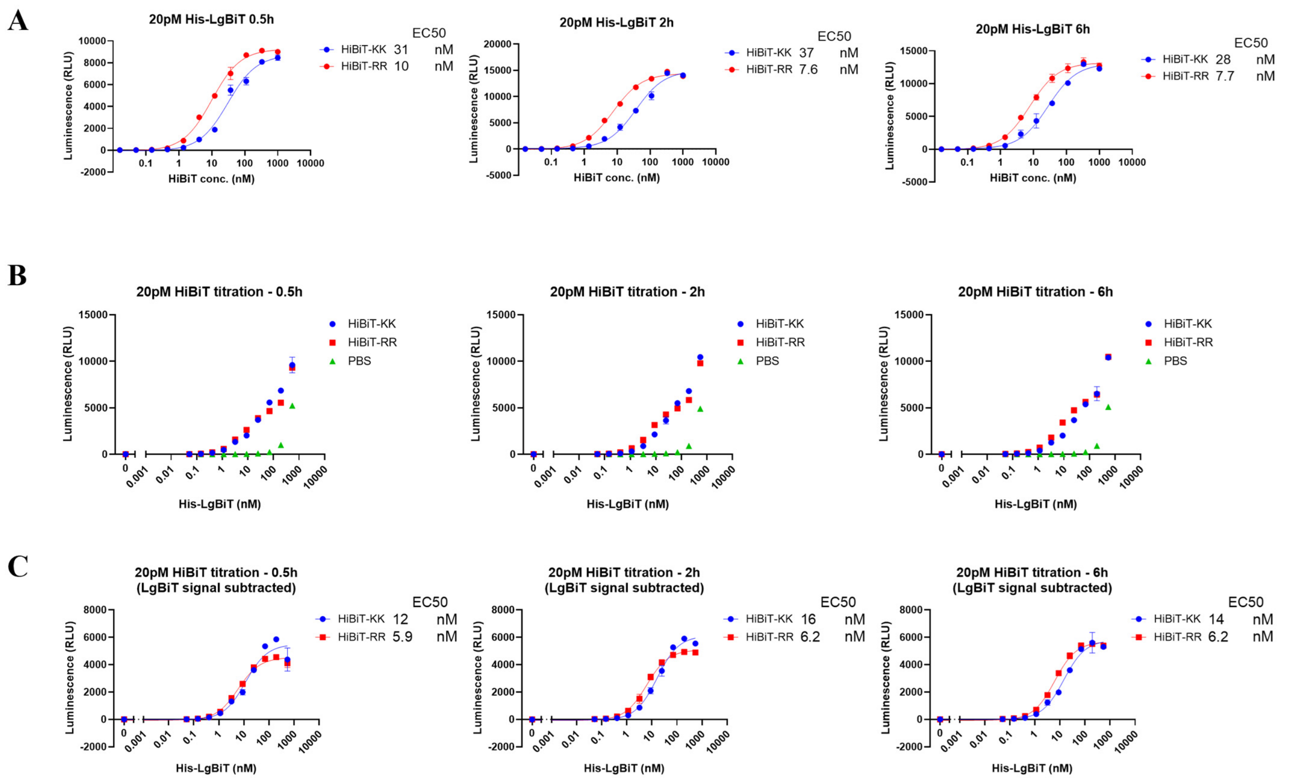


**Figure S1. HiBiT-RR showed comparable LgBiT protein binding affinity and similar luminescence output with the original HiBiT-KK. A.** The luminescence signal from 20 pM LgBiT protein when titrating with various concentrations of HiBiT-KK or HiBiT-RR for 0.5h, 2h and 6h. **B.** The luminescence signal from 20 pM HiBiT variants or PBS when titrating with various concentrations of His-LgBiT recombinant protein for 0.5h, 2h and 6h. **C.** The background LgBiT luminescence from PBS group were subtracted from panel B.

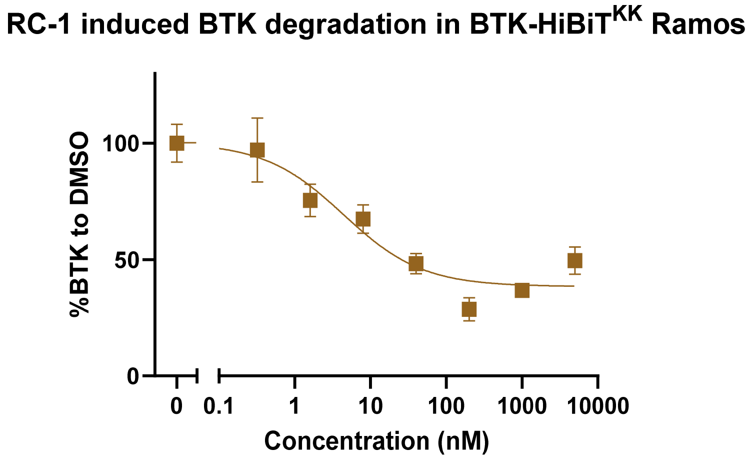


**Figure S2. RC-1 degradation potency of BTK in Ramos BTK(full length)-HiBiT$^{KK}$ knock-in cell line.** Cells were transferred into 96-well plate for 20,000 cells per well. Cells were incubated with DMSO or indicated compounds at 0.32, 1.6, 8, 40, 200, 1,000 nM, 5,000 nM in 1% DMSO for 24 hours. Bioluminescence signal was generated by adding furimazine substrate and LgBiT purified protein in the cell lysis buffer.

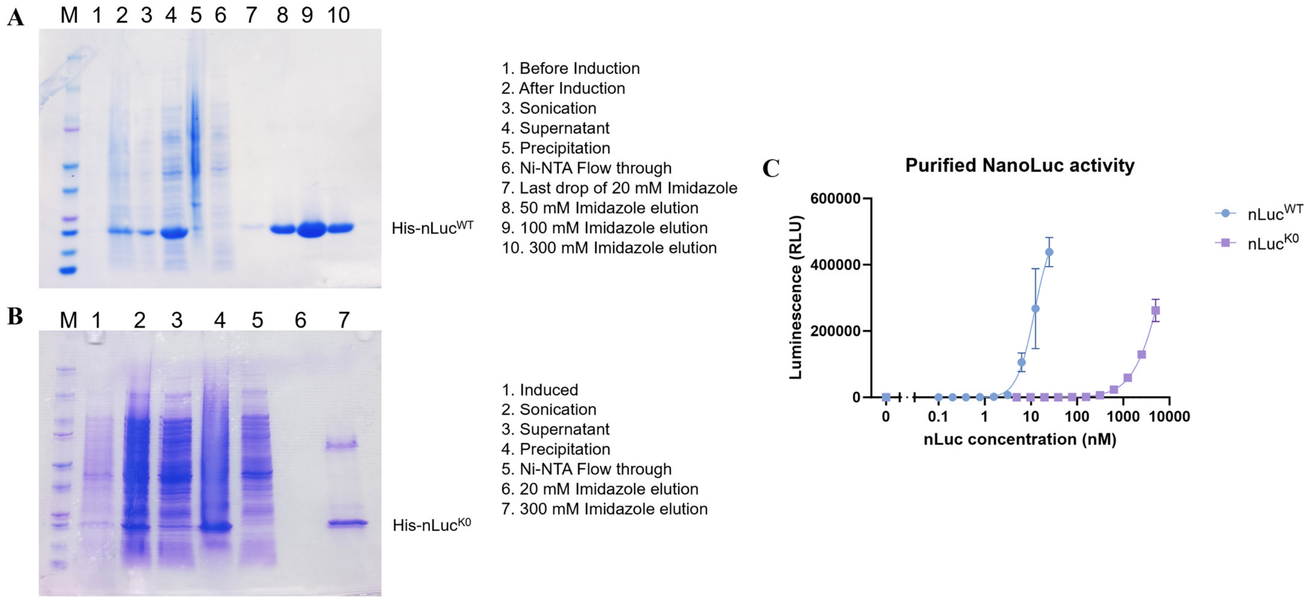


**Figure S3. A, B.** Protein purification of His-nLuc$^{WT}$ and His-nLuc$^{K0}$. **C.** Luciferase assay for purified nLuc WT and K0

**Table S1. RIPK1 degrader common structures**

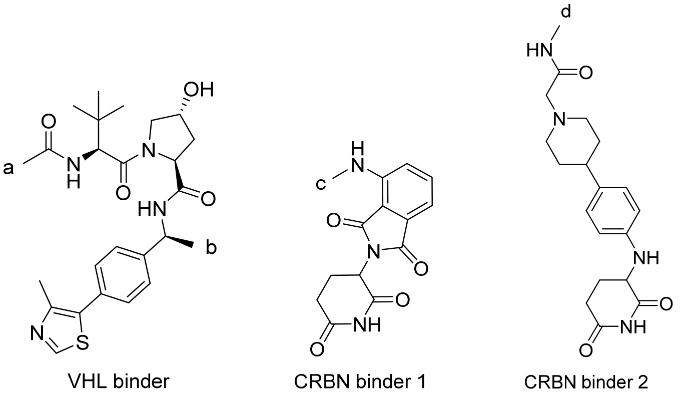


| Compound | E3 | Exit vector | Linker type |
|---|---|---|---|
| LD4172 | VHL binder | a | Flexible |
| LD4172NC | VHL binder (stereo-center at b flipped) | a | Flexible |
| LD5037 | VHL binder | a | Rigid |
| LD5042 | VHL binder | b | Flexible |
| LD5097 | VHL binder | b | Rigid |
| LD5063 | CRBN binder 1 | c | Flexible |
| LD5077 | CRBN binder 2 | d | Flexible |
| LD5084 | CRBN binder 2 | d | Rigid |
